# Selective motor stimulation of the pudendal nerve using multi-contact cuff electrodes: a pre-clinical study in feline and ovine models

**DOI:** 10.1101/2025.10.20.683457

**Authors:** Miguel Ortiz-Lopez, Amador C. Lagunas, Hera Akmal, Tim M. Bruns

## Abstract

**Introduction:** Pudendal nerve stimulation is a promising therapy for urinary incontinence, however stimulation can evoke off-target activity. We aimed to determine whether multi-contact cuff electrodes can selectively recruit motor fibers of the pudendal nerve trunk in preclinical feline and ovine models.

**Methods:** Multi-contact cuff electrodes were implanted around the pudendal nerve in anesthetized felines and ovines. Structured variations in electrode contact configurations and stimulation amplitudes were applied to evoke external urethral sphincter (EUS) and external anal sphincter (EAS) pressure responses. We calculated selectivity indices, EUS Scores, and EAS Scores to quantify selective recruitment and the magnitude of evoked pressure changes.

**Results:** We achieved selective motor activation, with preferential recruitment of the EUS or EAS in all three feline experiments and one of three ovine experiments. In felines, at least four electrode combinations selectively evoked EUS responses (EUS Score 0.5) and at least one combination targeted the EAS. In ovines, one EUS-selective and six EAS-selective combinations obtained comparable scores. In preliminary tests, we observed functional increases in leak point pressure and incontinence prevention with selective stimulation.

**Conclusions:** This study shows that multi-contact cuff electrodes can selectively activate EUS and EAS motor fibers in the pudendal nerve. Future work should focus on optimizing stimulation parameters to enhance selectivity and assess the translational potential of this approach for restoring pelvic organ control.

## Introduction

Stress urinary incontinence (SUI), defined as the involuntary leakage of urine during activities that increase abdominal pressure, significantly impacts quality of life and affects nearly one-third of women worldwide.^1,2^ Conventional therapies such as pelvic floor exercises or surgical interventions are often only partially effective, motivating the development of neurostimulation-based treatments that restore sphincter control.

The pudendal nerve, originating from sacral roots S2–S4, innervates the external urethral sphincter (EUS) and external anal sphincter (EAS) and provides sensory feedback from the external genitalia, urethra, and the perineal skin.^3^ Pudendal nerve stimulation (PNS) has emerged as a promising approach for restoring continence by activating motor pathways that reinforce urethral closure^4^. Current PNS methods that stimulate the pudendal trunk can produce variable motor recruitment^5^ and undesired stimulation effects^6^, which may reduce therapeutic precision. Achieving selective stimulation of pudendal trunk fibers could enhance sphincter control while minimizing off-target contractions and could also be leveraged to selectively engage pudendal sensory pathways for neuromodulation of lower urinary tract function. In clinical practice, selective stimulation may improve patient therapeutic outcomes by expanding the tolerable stimulation range and enabling effective responses at lower amplitudes.

Multi-contact nerve cuff electrodes offer a means to modulate current distribution along peripheral nerves and achieve selective activation of specific fascicles.^7^ Although computational modeling has suggested that this approach could enable selective pudendal stimulation^8^, its in vivo feasibility has not been validated. In this study, we evaluated whether multi-contact cuff electrodes can selectively recruit motor pathways within the pudendal nerve trunk. Using feline and ovine models, we compared EUS and EAS pressure responses across electrode configurations and stimulation amplitudes to assess the potential for selective sphincter control relevant to SUI therapy.

## Methods

### Animals

All procedures were approved by the local Institutional Animal Care and Use Committee (IACUC) and conducted in accordance with the NIH Guide for the Care and Use of Laboratory Animals. The experiments were carried out on five adult, purpose-bred, neurologically and sexually intact, female felines (0.95 ± 0.11 years old, 3.5 ± 0.2 kg at surgery) and five neurologically intact female ovines (41.1 ± 1.9 kg at surgery). Animals were obtained from a licensed commercial breeder (felines) and an approved agricultural supplier (ovines). Felines were housed in a 6.22 m × 2.9 m room with controlled temperature (19–21 °C), relative humidity (35–60%), a 12-hour light/dark cycle, ad libitum access to food and water, and environmental enrichment (toys and regular staff interaction). Ovines were group-housed in an open-air barn (12.2 m × 15.2 m) with continuous access to water and regular provision of feed.

The feline model was selected for its well-characterized pudendal nerve anatomy and established use in lower urinary tract studies.^9^ The ovine model was included to assess the stimulation approach in a larger animal with nerve dimensions more comparable to human anatomy and a history in bladder neuromodulation studies.^10,11^ Animals used in this study did not have SUI.

### Multicontact cuff electrode

For felines, a 20-micron Parylene C multi-contact cuff electrode (1.2-1.5 mm inner diameter, eight 0.70 mm^2^ electrode sites; Figure 1A, v1) was designed by our group and manufactured by the Polymer Implantable Electrode Foundry at the University of Southern California^12^, based on previously reported nerve dimensions.^13,14^ For ovines, the multi-contact cuff (1.5-2.0 mm inner diameter, eight 0.93 mm2 electrode sites; Figure 1A, v2) was modified to accommodate larger nerve sizes. In both designs, the electrode tip is rolled over the nerve to enclose the nerve and create the cuff electrode, with teeth on the tip locking into a slot to secure the cuff. Manufacturing and electrical characterization details are provided in the Supplementary Materials.

**Figure 1.**
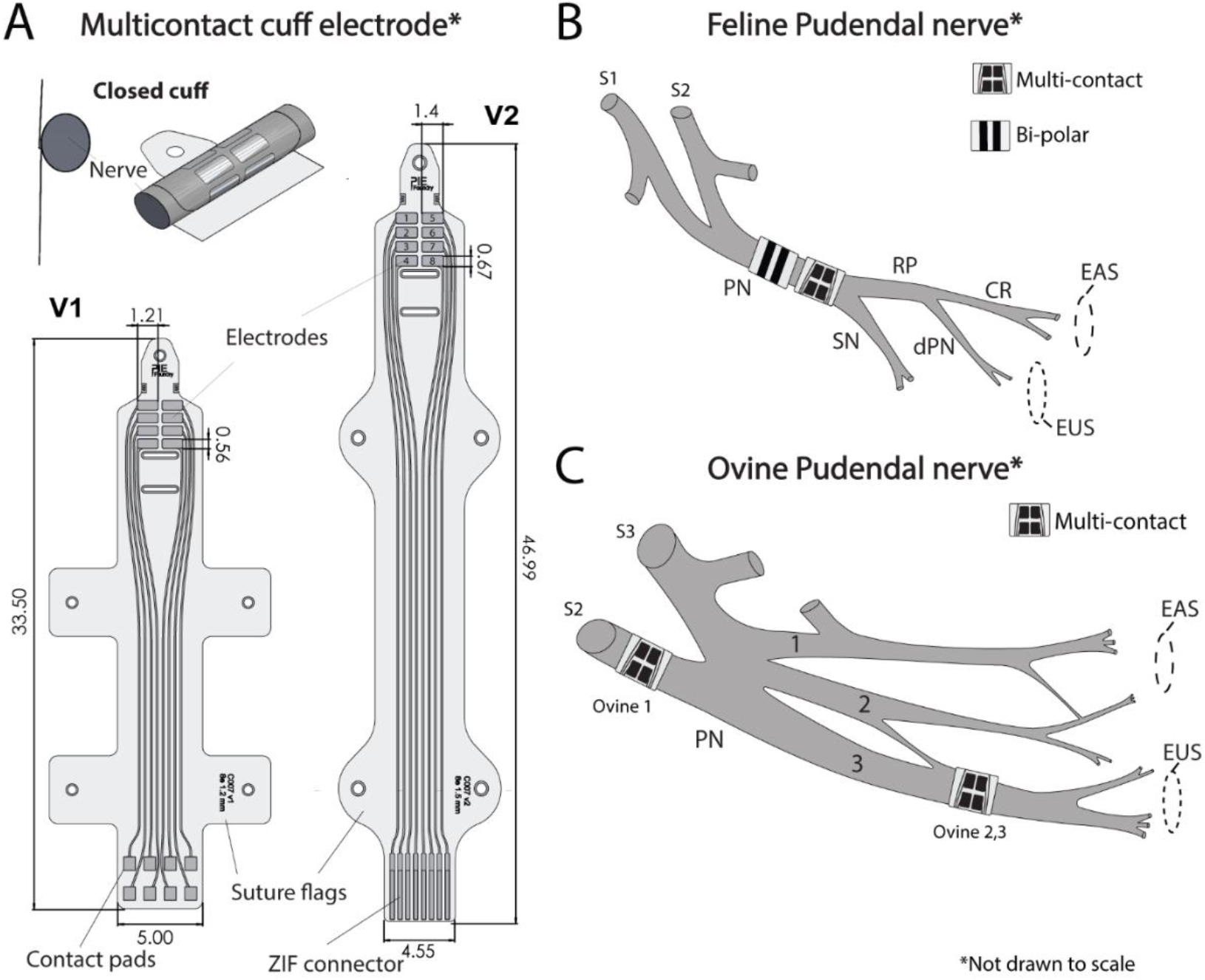
**A**. Multi-contact cuff electrode specifications. All measures are in mm. Electrode sites are numbered 1-8, as shown in the v2 cuff. **B**. Typical feline pudendal nerve anatomy within the ischiorectal fossa (lateral view) and planned implanted cuff electrode locations. Nomenclature follows Martin et al.^29^. Pudendal Nerve (PN), Rectal Perineal Nerve (RP), Sensory Nerve (SN). The PN trunk bifurcates into SN and RP, with the RP dividing into the Deep Perineal Nerve (dPN) and Caudal Rectal Nerve (CR). Both cuffs were planned to be placed at the PN trunk as shown in the figure; however, in Feline 2, the multi-contact cuff was placed at the RP. **C**. Illustrative ovine pudendal nerve anatomy based on our observations. A single pudendal trunk was not identified in any ovine. Instead, the pudendal nerve tends to be composed of three main branches; similar branching has been previously reported^30^. The actual branching pattern distal to the electrode site may be more complex than illustrated. EAS recruitment was observed for each electrode depicted here. In Ovine 1, we placed a multicontact cuff at a sacral nerve. In Ovine 2 and 3, we placed a multi-contact cuff on Branch 3 where electrical stimulation evoked the largest EUS response. (EAS – External Anal Sphincter, EUS – External Urethra Sphincter).

### Feline Surgical Procedure

Anesthesia was induced with a mixture of intramuscular Ketamine (6.6 mg/kg), Butorphanol (0.66 mg/kg), and Dexmedetomidine (0.011 mg/kg), followed by maintenance with 1-4% isoflurane. Heart rate, oxygen perfusion, temperature, and blood pressure were continuously monitored, and saline (10-20 ml/hr) was infused via a cephalic vein. A laparotomy was performed to place a 1-mm suprapubic Silastic catheter into the bladder dome via a small incision in the bladder wall and secured with a purse-string suture. In each animal, the left pudendal nerve was approached through the ischiorectal fossa using blunt dissection after a dorso-lateral incision, and a unilateral multi-contact cuff electrode was placed around the nerve (Figure 1A, v1). Nerve identity was confirmed by brief electrical stimulation to verify EUS and EAS responses before cuff placement. A bipolar cuff electrode (two mm inner diameter, 2-2.5 mm wire separation, 32 AWG-AS 636 Cooner Wire) was placed adjacent to the multi-contact cuff electrode for comparison testing. Additional catheters were placed transrectally for EAS (balloon, Laborie RPC-9) and transurethrally for EUS (3.5 Fr, Argyle) pressure measurement. Both catheters were positioned at the point of maximal sphincter pressure.

### Ovine Surgical Procedure

Ovines were given Xylazine (0.1-0.3 mg/kg) for catherization, and Propofol (2-6 mg/kg) was administered intravenously through the cephalic vein for induction, followed by 1-5% isoflurane maintenance. Heart rate, oxygen perfusion, temperature, and blood pressure were monitored, and an intravenous line was inserted in the cephalic vein for saline infusion at a rate of 10-20 ml/hr. After a dorso-lateral incision and reflection of the underlying muscle, the left sacral and pudendal nerve plexus was accessed for unilateral placement of the multi-contact cuff (Figure 1A, v2). Nerve identity was confirmed by brief electrical stimulation to verify EUS/EAS responses before cuff placement. The cuff was placed at a nerve location where both EUS and EAS pressure responses could be evoked, which varied among animals. A bipolar cuff electrode was not used for comparison testing in ovines due to reduced surgical and experimental time, as ovines are less hemodynamically stable under anesthesia as compared to felines.^15,16^ Catheters were placed trans-urethrally for bladder (3.5 Fr, Argyle) and EUS (solid-state pressure; 3.5 Fr Mikro-Cath, Millar), and trans-rectally for EAS (balloon; Laborie RPC-9) pressure measurement. Animals were euthanized with an intravenous overdose of pentobarbital (100 mg/kg).

### Experiment Setup and Data Collection

In felines, bladder, EUS, and EAS pressures were recorded via transducers (DPT-100, Utah Medical) and an amplifier (TBM4M, World Precision Instruments) with a PowerLab 8/35 system and LabChart 8 software (ADInstruments). In ovine, the EUS catheter was connected to an amplifier (FE224, ADInstrument). The multi-contact cuff electrode was routed through a custom relay system for automatic cycling through electrode combinations and pulse stimulator (Model 4100 Isolated Stimulator, A-M Systems), controlled via a custom MATLAB graphical user interface. The bipolar cuff electrode was connected directly to the stimulator. Symmetric biphasic rectangular pulses (210 µs, 30 Hz) were applied in bipolar mode. The EUS catheter was fixed at the urethra position with the maximal pressure response.

We applied a grid search across electrode combinations and up to three stimulation amplitudes per animal. In felines, amplitudes ranged from 75 to 500 μA; in ovines from 125 μA to 5 mA. All bipolar combinations using one electrode site per polarity were tested in every animal except Feline 1. In ovines, additional bipolar combinations comprising two electrode sites per polarity were also tested. Only electrode sites adjacent to each other were grouped (e.g., pairs 1-2 and 4-8, see Figure 1A v2). For each electrode combination, stimulation was applied for two seconds, with a 5-second interval between combinations. The three amplitudes were selected based on pressure responses observed during initial electrode testing for each animal. Trials with lower amplitudes were performed first.

As a preliminary proof-of-concept demonstration, urinary leakage prevention was evaluated in one feline and one ovine, using distinct protocols. EUS catheters were removed for this testing. In the feline, saline was infused until the bladder pressure leakage point, with and without PNS. Electrode combinations and amplitudes were selected based on evoked EAS responses to target both high and low off-target recruitment. Due to muscle fatigue during 30 Hz stimulation, stimulation was applied just before reaching the average no-stimulation bladder pressure leakage point and was stopped upon observation of leakage. The bladder was emptied and rested between trials. In the ovine, immediate urine leakage was observed upon infusion— a limitation previously documented in ovine models^10^, making pressure leakage point measurements uninformative. Consequently, we adopted a continuous saline infusion protocol: after constant leakage was established during infusion, stimulation was applied in five second on/off intervals with a different electrode combination used in each interval. Leaked urine was continuously measured using a weight scale, and each interval leakage rate was calculated in MATLAB.

### Data Analysis

We performed data preprocessing and analyses in MATLAB and statistical analyses in R. Each pressure recording was low-pass filtered with a cutoff frequency of 10 Hz. The pressure change was defined as the difference between the mean pressure during stimulation and the two-second baseline before stimulation, then normalized to the maximum multicontact cuff response per animal (*NP*). A selectivity index (*SI*) was calculated for each trial using the ratio between the normalized EUS response (*NP*_*Refer*_) and the normalized EUS and EAS (*NP*_*EAS*_) sum (1).

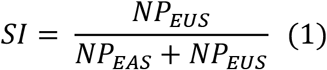

Since SI does not reflect the strength of muscle contractions, small EUS responses near the motor threshold can lead to a high SI for negligible EAS pressure changes. For trials where normalized pressure changes were below 0.2 for both the EUS and EAS, SI was set to 0.5. To combine selectivity and response magnitude, EUS and EAS Scores were computed per trial (2) and (3).

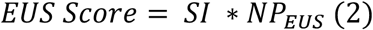

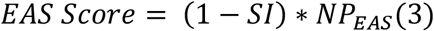

Scores greater than 0.5 indicate effective targeting of only one sphincter and at least 50% of its maximum pressure change. Kruskal-Wallis tests were performed to identify significant differences in EAS and EUS pressure responses and EUS and EAS Scores due to variations in stimulation amplitude, electrode configuration, and animal subject. Post-hoc Holm’s method was applied to adjust p-values for multiple comparisons. An adjusted p-value (p_adj_) less than 0.05 was considered significant.

Statistical comparisons for leakage testing were conducted separately for feline and ovine data. For the feline experiment, mean bladder pressure leakage points were compared between PNS and no-PNS conditions using a Mann-Whitney U test. For the ovine experiment, the mean urinary leakage rate was compared between electrode combinations that evoked low (< 5 cmH_2_O) and high (≥ 5 cmH_2_O) EUS pressure responses and intervals without stimulation, using a Kruskal-Wallis test, followed by a Dunn’s post hoc test. Significance was set at a p-value less than 0.05.

## Results

We performed surgical and experimental procedures on five felines and five ovines and characterized EUS and EAS responses in six animals. We encountered difficulties with two felines (one: urethral pressure catheter insertion failure; one: no stimulation response) and two ovines (no single nerve identified that evoked both EUS and EAS pressure changes simultaneously). The electrode implant locations are shown in Figure 1. In ovines, only the second slot of the multi-contact cuff (Figure 1A v2) was utilized due to the nerve size, resulting in partial coverage with the electrode contacts around the nerve.

Figure 2A illustrates examples of pressure traces obtained using different multi-contact or bipolar cuff electrode combinations in Feline 1. Figure 2B shows the distribution of Scores. For Feline 1, Feline 2, and Feline 3, a total of 16, 56, and 56 electrode combinations were tested for three different stimulation amplitudes, respectively. For each Ovine, 68 electrode combinations were tested for each stimulation amplitude.

**Figure 2.**
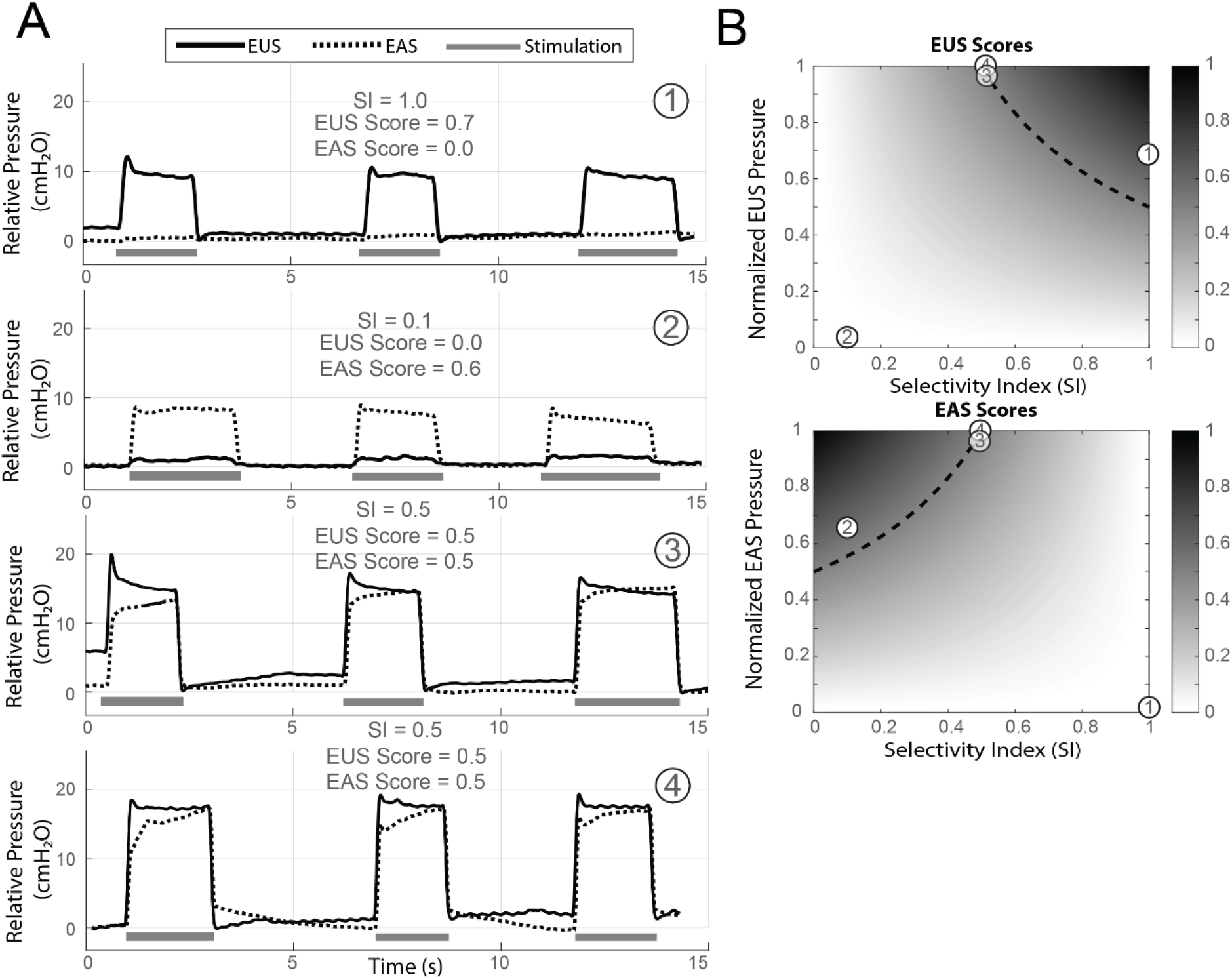
**A.** Examples of pressure traces for three different electrode combinations (Feline 1). Combination 1 (multi-contact cuff, SI = 1) shows exclusive EUS activation; Combination 2 (multi-contact cuff, SI = 0.1) shows predominant EAS activation; Combination 3 (multi-contact cuff, SI = 0.5) and Combination 4 (bipolar cuff, SI = 0.5) show balanced EUS–EAS activation. **B**. Distribution of EUS and EAS Scores. The dashed line represents the threshold where scores are equal to 0.5.

### Felines Evoked Pressure Responses, Selectivity, and Scores

Among all tested multi-contact cuff electrode combinations, *NP*_*EUS*_ values exceeded 0.5 in 10% of 48 electrode-amplitude combinations for Feline 1, 14% of 168 for Feline 2, and 18% of 168 for Feline 3 (Figure 3A). Six percent of the electrode combinations for Feline 1, 17% for Feline 2, and 24% for Feline 3 exhibited NP_EAS_ greater than 0.5 (Figure 3B). The median tendency of the SI across all animals and stimulation amplitudes hovered around 0.5 (Figure 3C). High EUS Score electrode combinations were observed across all felines, with 8% of the electrode combinations for Feline 1, 2% for Feline 2, and 4% for Feline 3 achieving EUS Scores greater than 0.5 (Figure 3D). Similarly, electrode combinations favoring the EAS were also observed across all felines, with 2% of Feline 1 trials, 8% of Feline 2 trials, and 10% of Feline 3 yielding EAS Scores greater than 0.5 (Figure 3E). The Kruskal-Wallis tests showed that stimulation amplitude (p_adj_ < 0.001), animal (p_adj_ < 0.001), and cathode location (padj < 0.05) significantly influenced EUS pressure changes. For EAS pressure changes, stimulation amplitude (p_adj_ < 0.001), animal (p_adj_ < 0.001), and the anode site (p_adj_ < 0.05) were significant. Both EUS and EAS Scores were significantly influenced by stimulation amplitude and animal (p_adj_ < 0.001 each); the cathode site influenced only EUS Score (p_adj_ < 0.01), while the anode site influenced only EAS Score (p_adj_ < 0.05).

**Figure 3.**
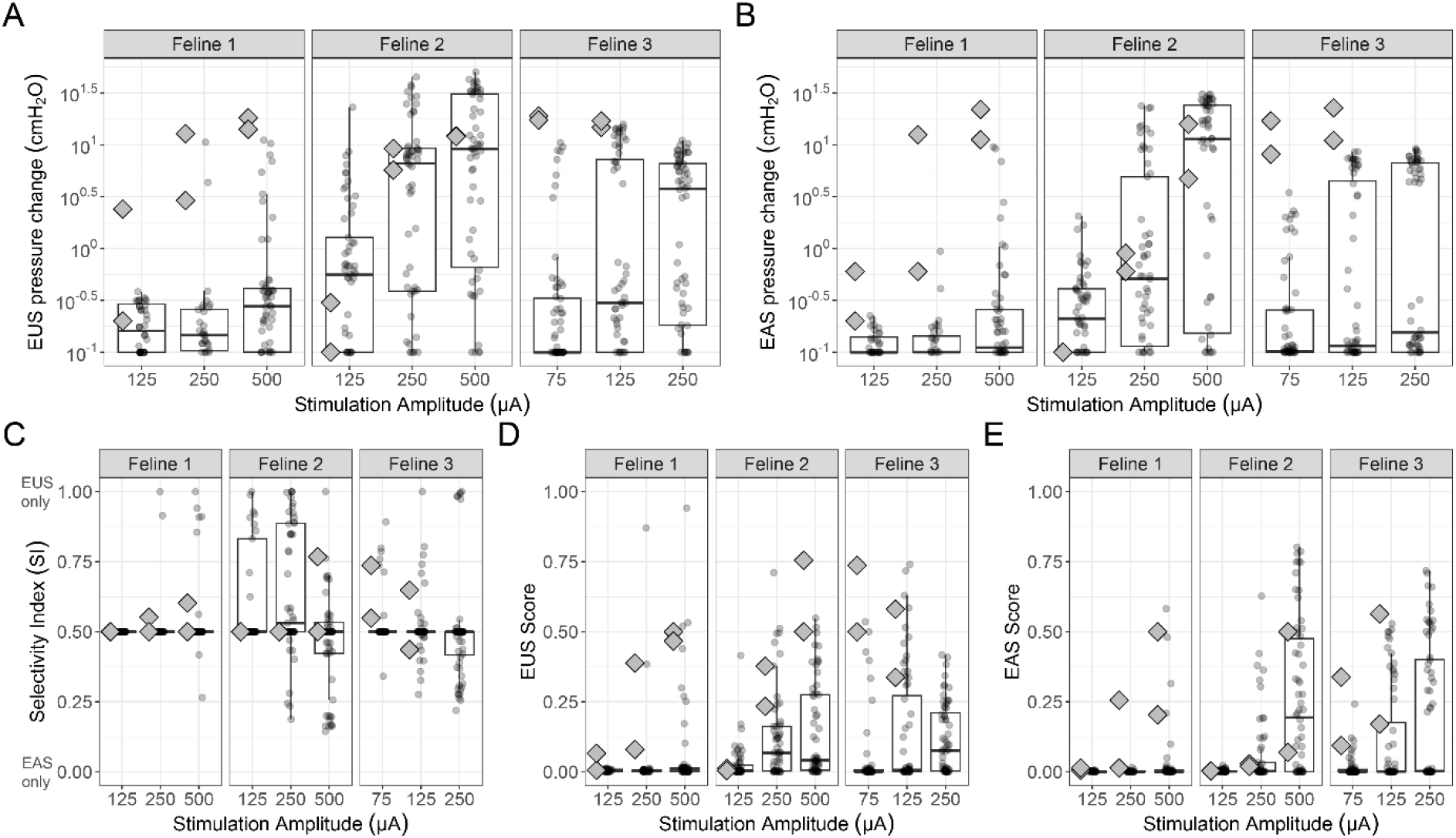
Summary results for feline experiments. **A.** Summary of EUS pressure changes. **B**. Summary of EAS pressure changes. A log transformation with an added offset of 0.1 cm H_2_O was applied to the pressure values for better illustration. **C-E**. Summary of Selectivity Indices, EUS Scores, and EAS Scores. In each figure, the box plots indicate the median, interquartile range (IQR), and the data range within 1.5 times the IQR with whiskers, for multi-contact stimulation. The diamonds indicate results for trials with a bipolar cuff electrode.

### Bipolar cuff vs Multicontact cuff

Among all trials with the bipolar cuff, 50% of six trials for Feline 1, 33% of six trials for Feline 2, and 100% of four trials for Feline 3 had *NP*_*EUS*_ values greater than 0.5. Similarly, 50% for Feline 1, 33% for Feline 2, and 100% for Feline 3 had *NP*_*EAS*_ values greater than 0.5. Only in Feline 2 did a multi-contact cuff electrode combination evoke larger EAS and EUS pressure changes than the bipolar cuff for all stimulation amplitudes — likely due to the more distal electrode placement (Figure 1B). At the same amplitudes, there were multi-contact cuff electrode combinations that achieved higher and lower SI than the bipolar cuff in all felines (Figure 1C). There was a multi-contact cuff combination that yielded the highest EUS Score (Figure 1D) and highest EAS Score (Figure 1E) for each feline, as compared to the bipolar cuff, except for the EUS score for Feline 2.

### Ovine Evoked Pressure Responses, Selectivity, and Scores

Among the tested combinations, 5% in Ovine 1 (of N=204 total electrode combinations), 18% in Ovine 2 (N=204), and 6% in Ovine 3 (N=204) had normalized mean EUS pressure change values greater than 0.5 (Figure 4A). Six percent of combinations in Ovine 1, 16% in Ovine 2, and 34% in Ovine 3 had normalized mean EAS pressure change values greater than 0.5 (Figure 4B). Only in Ovine 1 were the magnitudes of EUS and EAS responses similar, with much higher stimulation amplitudes required than in other ovines. The EUS pressure responses were approximately three times higher than the evoked EAS responses in Ovine 2 and 3, which was not unexpected due to the multi-contact cuff location on the nerve (Figure 1C). The median tendency across all animals and stimulation amplitudes hovered around 0.5 with a small IQR (Figure 4C). Ovine 2 exhibited electrode combinations with a wide range of SI, demonstrating both high SI (> 0.5) and low SI (< 0.5). In contrast, Ovine 1 and Ovine 3 showed primarily low SI. The EUS Scores varied among the Ovine, with only a small percentage achieving scores above 0.5: 0.5% in Ovine 1, 2.5% in Ovine 2, and 1% in Ovine 3. Kruskal-Wallis tests showed stimulation amplitude (p_adj_ < 0.001), animal (p_adj_ < 0.001), and cathode site (p_adj_ < 0.001) significantly affected EUS pressure changes. For EAS pressure, amplitude (padj < 0.001), animal (padj < 0.001), cathode site (p_adj_ < 0.001), and anode site (p_adj_ < 0.05) were significant. These factors also influenced EUS and EAS Scores (p_adj_ < 0.001), with anode site significant only for EAS Score (p_adj_ < 0.05).

**Figure 4.**
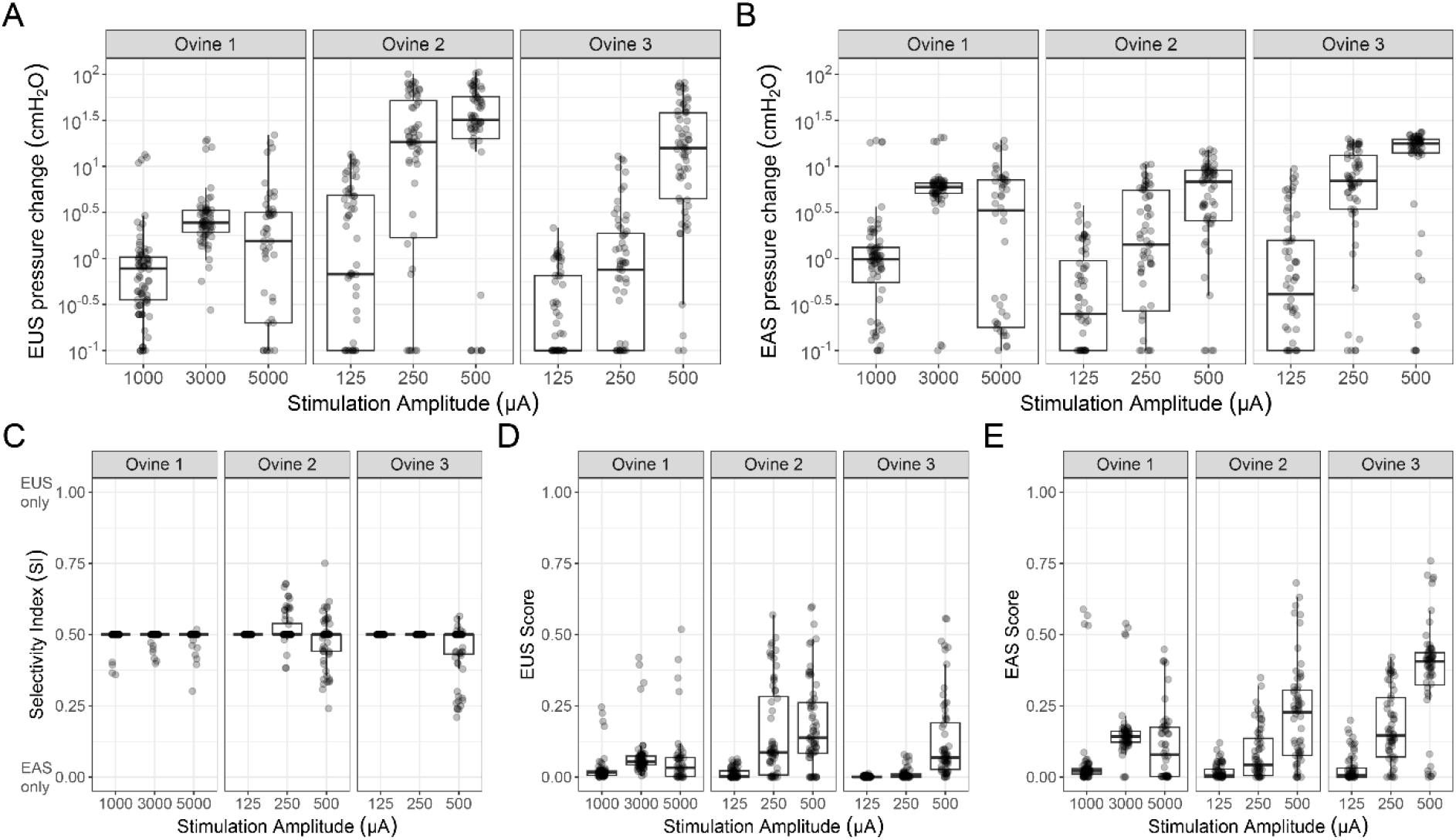
Summary results for ovine experiments. **A.** Summary of EUS pressure changes. **B**. Summary of EAS pressure changes. A log transformation with an added offset of 0.1 cm H_2_O was applied to the pressure values for better illustration. **C-E**. Summary of Selectivity Indices, EUS Scores, and EAS Scores. The box plots indicate the same statistics as in Figure 3.

### Urine Leakage Testing

In Feline 4 (Figure 5A), PNS significantly increased the bladder pressure leakage point (30.0 ± 5.1 cmH_2_O, n = 5) compared to bladder filling without stimulation (15.4 ± 2.5 cmH_2_O, n = 12; p-value < 0.001). Electrode combinations with low EAS responses (normalized values < 0.5) had a higher bladder pressure leakage point (35.4, 31.8, and 32.7 cmH_2_O, n=3) than those with high EAS responses (22.1 and 28.0 cmH_2_O, n =2). In Ovine 3 (Figure 6B), PNS reduced the rate of urinary leakage to an average rate of 9.3 ± 10.1 ml/min (n = 34) compared to intervals without stimulation (24.5 ± 8.9 ml/min, n = 35). Variations in stimulation-driven EUS responses were observed. Electrode combinations with higher EUS responses achieved a significantly lower rate of urinary leakage of (1.8 ± 3.6 ml/min, n = 16), compared to electrode combinations with lower EUS responses (16.0 ± 9.2 ml/min, n = 18; p-value <0.001) and no-stimulation (p-value <0.001).

**Figure 5.**
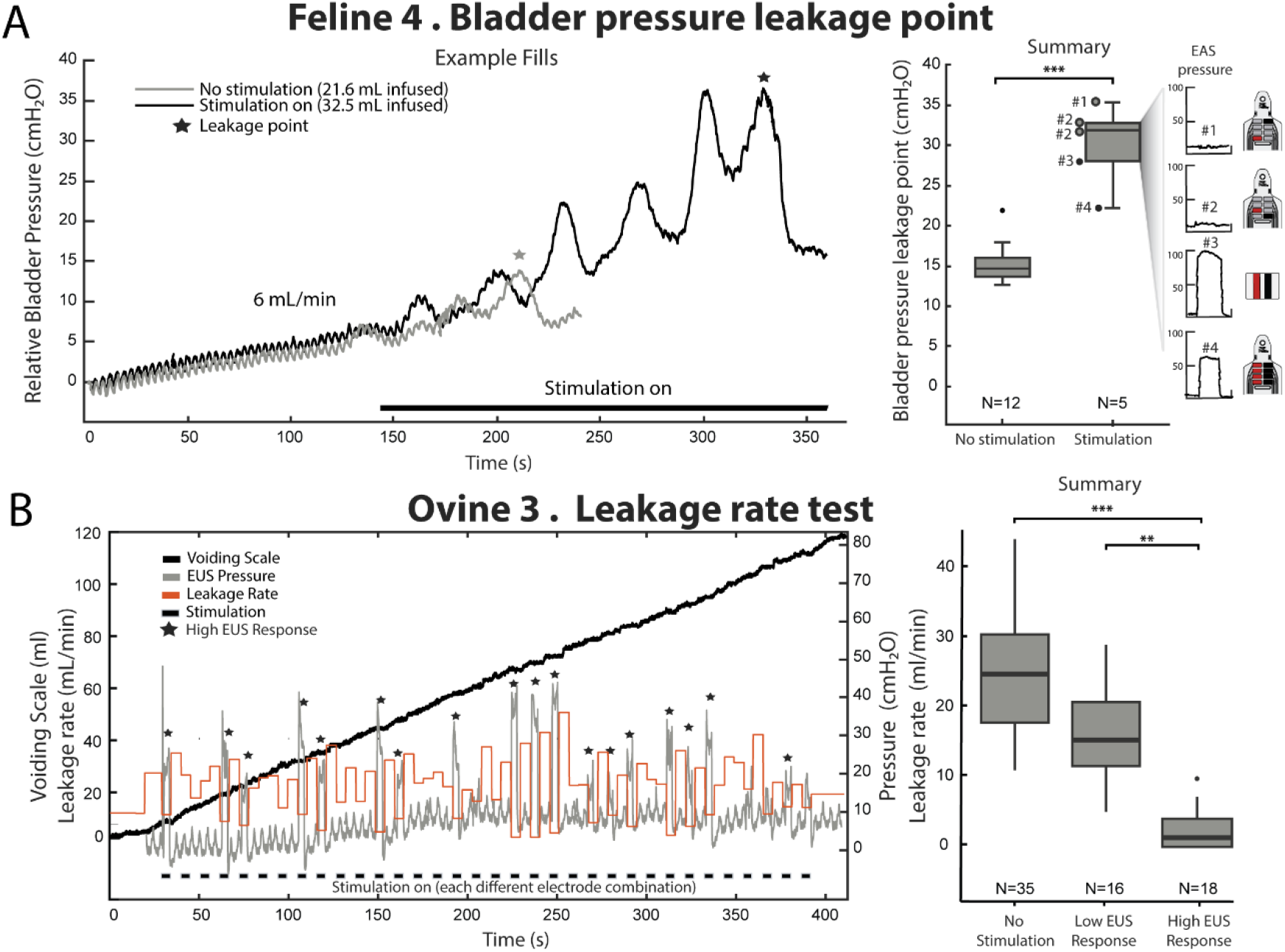
Urine leakage testing. **A.** Bladder pressure leakage point testing in Feline 4. Left: Representative example showing bladder filling sequences until leakage occurs. Right: Summary of leakage point bladder pressures during stimulation and no stimulation conditions. For the stimulation trials, the plots on the right show the external anal sphincter (EAS) pressure responses evoked during characterization trials for each electrode combination tested. **B**. Leakage rate testing in Ovine 3. Left: Voiding sequence, illustrating the leakage rate during on-off sequences of PNS. The overlaid EUS pressure was measured during a separate test with a catheter in place, using a matched stimulation-electrode ordering and timing. Right: Summary of leakage rates during periods without stimulation and electrode combinations that evoked low EUS responses (<5 cmH_2_O) or high EUS responses (≥5 cmH_2_O). (***p-value < 0.001, **p-value < 0.01).

## Discussion

### Evoked Pressure Responses and Nerve Anatomy Differences

The primary factors influencing EUS and EAS pressure responses were stimulation amplitude and animal subject, which likely reflect electrode placement location and nerve anatomy differences (Figure 1B). The anatomical organization of the pudendal nerve varied notably between felines and ovines. In ovines, the absence of a single pudendal nerve trunk and the limited anatomical literature for this species made experimental planning more challenging compared to felines. Substantial anatomical variation has also been documented in the human pudendal nerve, including differences in trunk formation and branching patterns^17,18^, suggesting that similar variability will need to be considered when translating selective pudendal trunk stimulation strategies into clinical practice. Higher stimulation amplitudes generally yielded higher responses for both the EUS and the EAS; except Ovine 1, response fatigue was observed at the highest stimulation amplitude. Notably, Ovine 1 required a larger stimulation amplitude to evoke responses congruent with the placement of the multi-contact along the nerve. Despite these differences, the dynamics of EUS-evoked pressure responses — specifically, the rapid onset and subsequent plateau — were consistent across species and similar to those observed in canines and felines^19,20^.

### Electrodes Combination Effect on Selectivity and Scores

EUS and EAS selective activation was more consistently achieved in felines than in ovines. We observed a wide range of SI by varying electrode combinations and stimulation amplitudes in felines. In ovines, selectivity was generally low, with EUS and EAS often co-activated. Similarly, EAS and EUS scores were higher in felines. These observations of inter-subject variability in ovines may be attributed to the differences in the electrode placement, limited circumferential coverage by the multi-contact cuff, larger nerve size, and fascicular organization. This variability reflects clinical observations that PNS using electrode leads that lack full circumferential coverage resulted in variable motor responses between individuals^5^.

In both animal models, stimulation amplitude was the main factor driving EUS and EAS Scores, however electrode combinations also significantly influenced outcomes at effective stimulation levels. Notably, the cathode electrode site had a greater effect on EUS Scores than the anode site, while the anode site affected EAS Scores. The cathode typically causes depolarization, which affects nearby fibers, while the anode shapes the electric field and influences which fibers are activated.^21^ Our results suggest that for these animal and cuff locations, fibers innervating the EUS are likely closer to the nerve surface and thus more affected by the localized depolarization caused by the cathode, while fibers controlling the EAS may lie deeper or more centrally within the nerve, influenced more by the broader electric field shaped by the anode. This interpretation is supported by the observation that higher EUS scores were achieved at lower stimulation amplitudes than EAS Scores, with correspondingly lower motor thresholds for the EUS. However, this interpretation is inferred from selectivity patterns rather than direct anatomical mapping and should be confirmed in future studies with histological correlation.

These findings are highly specific to our particular electrode placements and animal subjects. Consequently, electrode placement variations or different nerve fascicle organization might lead to different outcomes. Our findings align with the insights from Kent and Grill, who demonstrated computationally that electrode positioning around the pudendal nerve and fascicle organization significantly influences stimulation selectivity^8^. Despite the challenges posed by varying anatomies and electrode placements, our findings support the potential of multi-contact cuffs for selective PNS.

### Bipolar cuff versus Multicontact cuff

Our data indicate that multi-contact cuffs can achieve higher selectivity indices at the same stimulation amplitudes and higher EUS Scores at lower stimulation amplitudes compared to bipolar cuffs. Lower stimulation amplitudes reduce the likelihood of stimulation-related discomfort and adverse events such as nerve damage, habituation, and muscle fatigue in clinical settings.^22^ Although the bipolar cuff evoked larger pressure responses than the multi-contact cuff, due to their different electric fields, unintended activation is more likely. Multi-contact cuffs allow better control of an electric field’s size and direction.

### Urine Leakage Testing

Our pilot results demonstrate the potential of multi-contact PNS for preventing urinary leakage. In the feline experiment, selective PNS effectively increased the bladder pressure leak point (Figure 5A). This is consistent with a previous feline study that reported that PNS can significantly increase leak point pressure^23^. In the ovine model, we identified electrode combinations that enhanced EUS responses and improved leakage control (Figure 5B). Collectively, these results show the potential of selective PNS using multicontact cuffs for preventing or reducing SUI. These results were obtained from only two neurologically intact animals. Future studies should evaluate efficacy in SUI models with appropriately powered cohorts

### Limitations, Clinical Perspectives, and Future Directions

Our study was limited to a relatively small sample size of six animals. Larger cohorts and refined methodologies would confirm and expand upon our findings. We acknowledge the inherent differences between animal models and human physiology, and the anatomically diverse structure of the pudendal nerve in humans.^17,24^ While our results from animal models are promising, translating these findings to human applications requires a careful comparison of fascicle organization to ensure relevance and applicability. Another limitation of this study is that we used pressure measurements rather than EMG recordings, which cannot unambiguously distinguish between direct motor activation and reflex-mediated contraction. However, the rapid onset and stimulus-locked timing of observed pressure responses, combined with the use of isoflurane anesthesia (which suppresses spinal reflexes), suggest that direct motor activation was the primary mechanism for observed pressure responses.

Our experimental design did not identify optimal stimulation amplitudes for each electrode combination. Previous studies with multi-contact cuffs for other applications have shown that each electrode combination has a distinct selectivity window,^25–27^ so optimizing parameters per combination would likely improve selectivity scores. Additionally, key stimulation parameters, such as pulse width and frequency, were held constant during testing because exhaustive exploration becomes impractical due to the exponential growth in possible parameter combinations. Efficient methods are needed to navigate this complex parameter space and enable clinical translation. Also, the clinical application of pudendal nerve cuff electrodes faces challenges due to the invasiveness of electrode placement compared to current practices. Advances in surgical techniques may facilitate cuff placement.^28^Future studies should evaluate selective PNS efficacy in SUI models and directly compare multi-contact and conventional electrode approaches for functional incontinence prevention.

## Conclusion

This study demonstrates the potential of multi-contact cuff electrodes for selective PNS. Despite variability among the animal subjects, our findings indicate that multi-contact electrodes can achieve selective motor responses more efficiently than standard bipolar cuffs at lower stimulation amplitudes. Multi-contact cuff electrodes may improve the management of urinary and fecal incontinence by specifically targeting efferent fibers while minimizing off-target effects. Future research should refine electrode designs and develop efficient stimulation parameter optimization methods.

## Acknowledgments

The multi-contact cuff electrodes described in this report were provided by the Polymer Implantable Electrode Foundry at the University of Southern California and funded under the BRAIN Award (U24NS113647). The authors thank Eric Kennedy, Brinton Vandegrift, and Mark Nissen for their assistance with data collection. We also thank Samuel Shrift, Kee Scholten, and Paras Patel for their contributions to the design and evaluation of the multi-contact cuff electrodes. Additionally, we extend our gratitude to the University of Michigan Unit for Laboratory Animal Medicine for their assistance with animal care.

