## Supplemental for "Selective motor stimulation of the pudendal nerve using multi-contact cuff electrodes: a pre-clinical study in feline and ovine models"

### Supporting Information

#### A. Multi-contact cuff electrodes

Details about the manufacturing process can be found in Scholten et al. [1] In version 1, Wires (76.2 µm platinum iridium bare, 139.7 µm perfluoroalkoxy alkanes (PFA) coated, A-M Systems, Sequim, WA, USA) were soldered to the cuff electrode contact pads using silver epoxy (H20E, Epoxy Technology, Inc., Billerica, MA, USA). After the silver epoxy was cured, insulation epoxy (353ND-T, Epoxy Technology, Inc., Billerica, MA, USA) was applied over the exposed wires on the contact pads. In the second version, the contact pads were modified to match a zero-force connector (WR-FPC 0.50 mm SMT ZIF Horizontal, Würth Elektronik eiSos GmbH & Co. KG, Germany).

#### B. Electrochemical impedance spectroscopy (EIS) and cyclic voltammetry (CV) for multi-contact cuff electrodes

EIS and CV measurements were taken with a PGSTAT12 Autolab potentiostat (Metreohm/Eco Chemie, Utrecht, Netherlands) controlled by the vendor-supplied NOVA software. EIS and CV measurements were analyzed using custom Matlab (Mathworks, Natick, MA) scripts. The electrodes sites were submerged by 1 mm in 1x phosphate-buffered saline (PBS, BP3994, Fisher, Waltham, MA). An Ag|AgCl electrode (RE-5B, BASi, West Lafayette, MA) served as a reference electrode, and a stainless steel rod was selected as the counter. EIS measurements were obtained by applying a 10 µVRMS signal from 10Hz to 31 kHz. Impedance magnitude and phase data were extracted for each electrode site. Electrode sites with impedance values above 5 kΩ at 1 kHz were excluded from further analysis. For all remaining electrode sites, the mean and standard deviation of impedance magnitude and phase were computed as a function of frequency, and the average impedance at 1 kHz was calculated. CV measurements were performed by sweeping the potential from 0.8 V to −0.6 V and back to 0.8 V at a scan rate of 1 V/s, for three consecutive cycles. For each electrode site, current signals were extracted, aligned, and averaged across repeated scans. The full charge storage capacity was calculated by integrating the current over the entire CV potential range and normalizing to electrode surface area (µC/cm²). One electrode contact used in

Feline 1 was non-functional. The average electrode contact impedances at 1 kHz for multicontact cuffs v1 and v2 were 0.93 ± 0.16 kOhms (n = 23) and 1.0 ± 0.18 kOhms (n = 24), respectively. The full charge capacities, calculated as the time integral of the absolute current across the voltage window and normalized to the contact area, were 1323 ± 174 μC/cm² for v1 and 1255 ± 81 μC/cm² for v2. Figures providing EIS and CV data across all cuff electrode contacts are given in Figure S1.

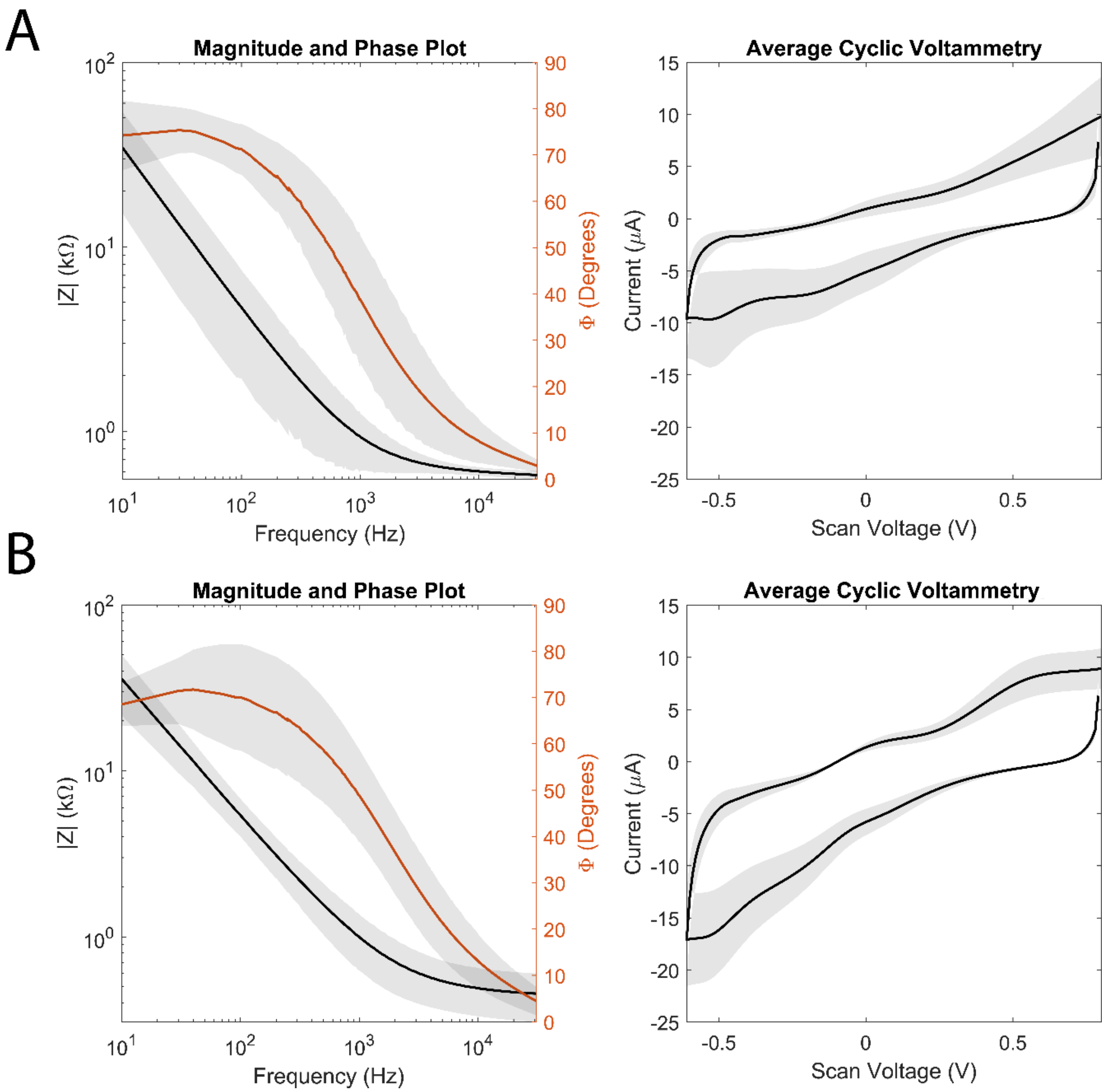


**Figure S1**. Electrochemical Impedance spectroscopy (EIS) and cyclic voltammetry (CV) across A. Version 1 multi-contact cuff electrodes and B. Version 2 electrodes. EIS data are presented as logarithmic plots of average impedance magnitude versus frequency with overlaid phase. Shaded regions represent ±2 standard deviations. The average electrode contact impedances at 1 kHz for multicontact cuffs v1 and v2 were 0.93 ± 0.16 kOhms (N = 23 electrode sites) and 1.0 ± 0.18

kOhms (N= 24), respectively. CV data are shown as averaged cyclic voltammograms, with shaded error bands indicating ±2 standard deviations across electrodes.

**Table S1.** Summary of Kruskal-Wallis Test Results for Pressure Measures and Score Indices in Feline and Ovine Multicontact-Cuff Stimulation Experiments. $\chi^2$ and adjusted p-values are reported for each variable and outcome measure.

| Species | Measure | Variable | $\chi^2$ | Adjusted p-value |
|---|---|---|---|---|
| **Feline** | **EUS Pressure** | Stimulation amplitude | 23.6 | 9e-5 |
| | | Feline | 58.1 | 1e-12 |
| | | Cathode site | 17.12 | 0.033 |
| | | Anode site | 9.38 | 0.2 |
| | **EAS Pressure** | Stimulation amplitude | 23.6 | 9e-5 |
| | | Feline | 49.9 | 5e-11 |
| | | Cathode site | 13.2 | 0.067 |
| | | Anode site | 18.1 | 0.02 |
| | **EUS Score** | Stimulation amplitude | 39.0 | 6e-8 |
| | | Feline | 17.8 | 4e-4 |
| | | Cathode site | 22.9 | 0.0035 |
| | | Anode site | 9.87 | 0.196 |
| | **EAS Score** | Stimulation amplitude | 17.6 | 0.0016 |
| | | Feline | 23.48 | 3e-5 |
| | | Cathode site | 13.79 | 0.055 |
| | | Anode site | 20.06 | 0.018 |
| **Ovine** | **EUS Pressure** | Stimulation amplitude | 172.79 | 7e-35 |
| | | Ovine | 44.34 | 5e-10 |
| | | Cathode site | 88.3 | 8e-11 |
| | | Anode site | 23.06 | 0.181 |
| | **EAS Pressure** | Stimulation amplitude | 177.9 | 6e-36 |
| | | Ovine | 31.4 | 3e-7 |
| | | Cathode site | 92.7 | 1e-11 |
| | | Anode site | 30.3 | 0.034 |
| | **EUS Score** | Stimulation amplitude | 179.8 | 2e-36 |
| | | Ovine | 38.8 | 7e-9 |
| | | Cathode site | 78.2 | 5e-9 |
| | | Anode site | 38.8 | 0.077 |
| | **EAS Score** | Stimulation amplitude | 186.57 | 8e-38 |
| | | Ovine | 20.97 | 6e-5 |
| | | Cathode site | 82.81 | 6e-8 |
| | | Anode site | 29.6 | 0.04 |
